# Opponent processing in the retinal mosaic of nymphalid butterflies

**DOI:** 10.1101/2022.02.10.479932

**Authors:** Primož Pirih, Marko Ilić, Andrej Meglič, Gregor Belušič

## Abstract

The eyes of nymphalid butterflies, investigated with incident illumination, show colourful facet reflection patterns, the eye shine, which is uniform or heterogeneous, dependent on the species. Facet colours suggest that the ommatidia contain different sets of photoreceptors and screening pigments, but how the colours and the cell characteristics are associated has not been clearly established. Here we analyse the retinae of two nymphalids, *Apatura ilia*, which has a uniform eyeshine, and *Charaxes jasius*, a species with a heterogeneous eye shine, using single-cell recordings, spectroscopy and optical pupillometry. *Apatura* has UV, blue and green-sensitive photoreceptors, allocated into three ommatidial types. The UV and blue-sensitive cells are long visual fibres (LVFs), receiving opponent input from the green-sensitive short visual fibres (SVFs). *Charaxes* has an expanded set of photoreceptors, allocated into three additional, red-reflecting ommatidial types. All red ommatidia contain green-sensitive LVFs, receiving opponent input from red receptors. In both species, the SVFs do not receive any opponent input. The simple retina of *Apatura* with three ommatidial types and two colour-opponent channels can support trichromatic vision. *Charaxes* has six ommatidial types and three colour-opponent channels. Its expanded receptor set can support tetrachromatic vision.

## I. Introduction

Colour vision is based upon the analysis of the spectral composition of the visual scene. The neural signals originating from photoreceptors with different spectral sensitivities are compared in a process of colour opponency. The spectral properties of colour opponent visual neurons match the statistics of natural scenes, maximizing the signal variance and allowing for optimal information processing in the visual pathway [1,2]. The optimal transformation for trichromatic vision is achieved through decomposing the retinal signals into an achromatic channel and two colour-opponent channels [3]. In flies and butterflies, the first stage of colour processing is performed already by the photoreceptors that directly inhibit each other through inter-photoreceptor synapses with histaminergic chloride channels [4,5,6,7,8]. Colour processing is continued downstream in the optical ganglia, the lamina and the medulla [5,8,9]. Here, we study colour opponency in the retinal mosaic of brush-footed butterflies (Nymphalidae).

Compound eyes are built from discrete optical units, the ommatidia. In all butterfly families with afocal apposition eyes studied thus far (families Papilionidae, Pieridae, Lycaenidae, Nymphalidae), each ommatidium contains eight large photoreceptors R1-8 and a small basal R9. The photoreceptors contribute their light sensing microvilli to the common light guide, the rhabdom. The microvilli of R1,2 are oriented vertically (i.e. parallel to the dorso-ventral eye axis), horizontally in R3,4, and diagonally in R5,7 & R6,8. The horizontal and diagonal photoreceptors R3-4 & R5-8 are the short visual fibres (SVF) that are presynaptic to the large monopolar neurons of the lamina [10,11], which in turn condition the signals for processing of achromatic contrasts and motion. In most butterflies studied so far, the SVFs are expressing a long-wavelength (LW) rhodopsin, peaking in the green range (520∼560 nm), but exceptions do exist: in the dorsal eye of the lycaenid butterfly *Lycaena rubidus*, blue opsin mRNA is co-expressed with LW opsin mRNA in the females, and exclusively expressed in the males [12].

The vertical photoreceptors R1&2 and the basal R9 are the long visual fibres (LVF) that project axons directly to the medulla and contribute to the processing of colour and polarisation content of the visual scene. In most studied butterflies, receptors R1&2 express the short-wavelength (SW), ultraviolet (U, 340∼370 nm) or blue-peaking (B, 420∼460 nm) opsins [13,14]. The U and B receptors allocated to R1-2 give rise to three ommatidial {UU, UB, BB} that are randomly distributed across the retina with species-specific fractions [15]. The random expression of the various visual pigments is driven by the *spineless* mechanism [16]. The basal cells R9 likely express the same LW opsin as R3-8 [13].

The retinae of papilionid and pierid butterflies are fully tiered [17]. The rhabdomeres of R1-4 occupy the distal tier of the rhabdom and the rhabdomeres R5-8 constitute the proximal tier. The SW part of the downwelling light is filtered by the visual and screening pigments in the distal tier, resulting in red-shifted sensitivities of the proximal photoreceptors R5-8 [18,19]. In papilionids, the basal LVF R9 can be either green- or red-sensitive [20] and expresses the LW opsin mRNA [21]. The red-sensitive basal LVF photoreceptor in pierids [11] likely also expresses a LW opsin. The inter-photoreceptor opponent synapses are located both downstream in the optic ganglia – the lamina and the medulla – and in the retina, along the photoreceptor axons, rendering the opponent signals detectable by intracellular recordings from the photoreceptors [5,6,7]. In the Japanese swallowtail, *Papilio xuthus*, opponent signals were detected in all photoreceptors R1-8 [6].

In brush-footed butterflies (nymphalids), the retina is incompletely tiered. The rhabdomeres of R3-8 are present along the entire length of the ommatidium, while the rhabdomeres of R1&2 may be present only distally [10,22] or may span along the whole rhabdom [13,23]. A tracheolar basket at the base of the retina forms the reflective *tapetum lucidum* in all studied butterfly families with apposition eyes [24,25] except in the papilionids [26]. The light launched into the rhabdom is reflected from the tapetum, bringing about *the eye shine*. The reflections from individual ommatidia are coloured, depending on the composition of visual pigments (rhodopsin R and its isomere metarhodopsin M) and the presence of screening pigments. A red ommatidial colour is a tell-tale sign for red (i.e. blue and green-absorbing) screening pigment being apposed to the rhabdom. Ommatidia without such a screening pigment appear yellow, green, pale or blue, depending on the tuning of the tapetum, rhodopsin composition and rhabdom length [15,25,27,28]. In species with a uniform eye shine, e.g. *Vanessa atalanta*, vertical cells R1&2 express exclusively UV or blue-peaking rhodopsins, similarly as in papilionids and pierids [13,28].

Many nymphalids have a non-uniform eye shine with red-reflecting ommatidia [25,29], which may indicate the presence of more than three ommatidial types. For instance, in some *Heliconius* butterflies, R1&2 can express another type of UV opsin (UV2, due to opsin duplication) [14] or a green-absorbing (G; long wavelength, LW) opsin [30], which leads to the expansion of possible ommatidial types and a complex retinal mosaic. Those R1&2 that express LW opsin likely reside in the red ommatidia, which furthermore contain a functional, red-sensitive R9 that inhibits the green-sensitive vertical cells [7]. The functional significance of R9 in the non-red ommatidia is currently unknown, but their spectral sensitivity is likely less red shifted [15,28], similarly as in *Papilio* [20].

Here, we study the inter-photoreceptor opponent mechanisms in brush-footed butterflies (family Nymphalidae). We focus our study on the lesser purple emperor *Apatura ilia*, and the two-tailed pasha, *Charaxes jasius*, the former with a simple and the latter with a complex eye mosaic. We show that the emperor is equipped with the basic trichromatic photoreceptor set, while the pasha has additional five photoreceptor types. We show that the expanded set of receptors in the pasha (green-sensitive R1&2, yellow-sensitive R3-9 and red-sensitive R9) is allocated to the red ommatidia. We identify two and three colour opponent channels in the emperor and the pasha, respectively. Taken together, the eyes of brush-footed butterflies can have either a simple retinal mosaic with three ommatidial types and two colour opponency channels, or a complex retinal mosaic with six ommatidial types and three opponency channels that are presumably used as the substrate for tetrachromatic vision.

## II. Results

### (a) Eyeshine and rhodopsin photochemistry

The eye shine mosaic in *Charaxes jasius* is predominantly green in the dorsal part and yellow-green in the central and ventral part, speckled with red ommatidia whose fraction is higher ventrally (figure 1*a* *below*). The red ommatidia stand out prominently in the hyper-spectral image of the dark-adapted eye of *Charaxes* (figure 1*f* *below*), while in the eye of *Apatura*, mosaic regionalisation is not observed (figure 1*g,j* *below*).

**Figure 1.**
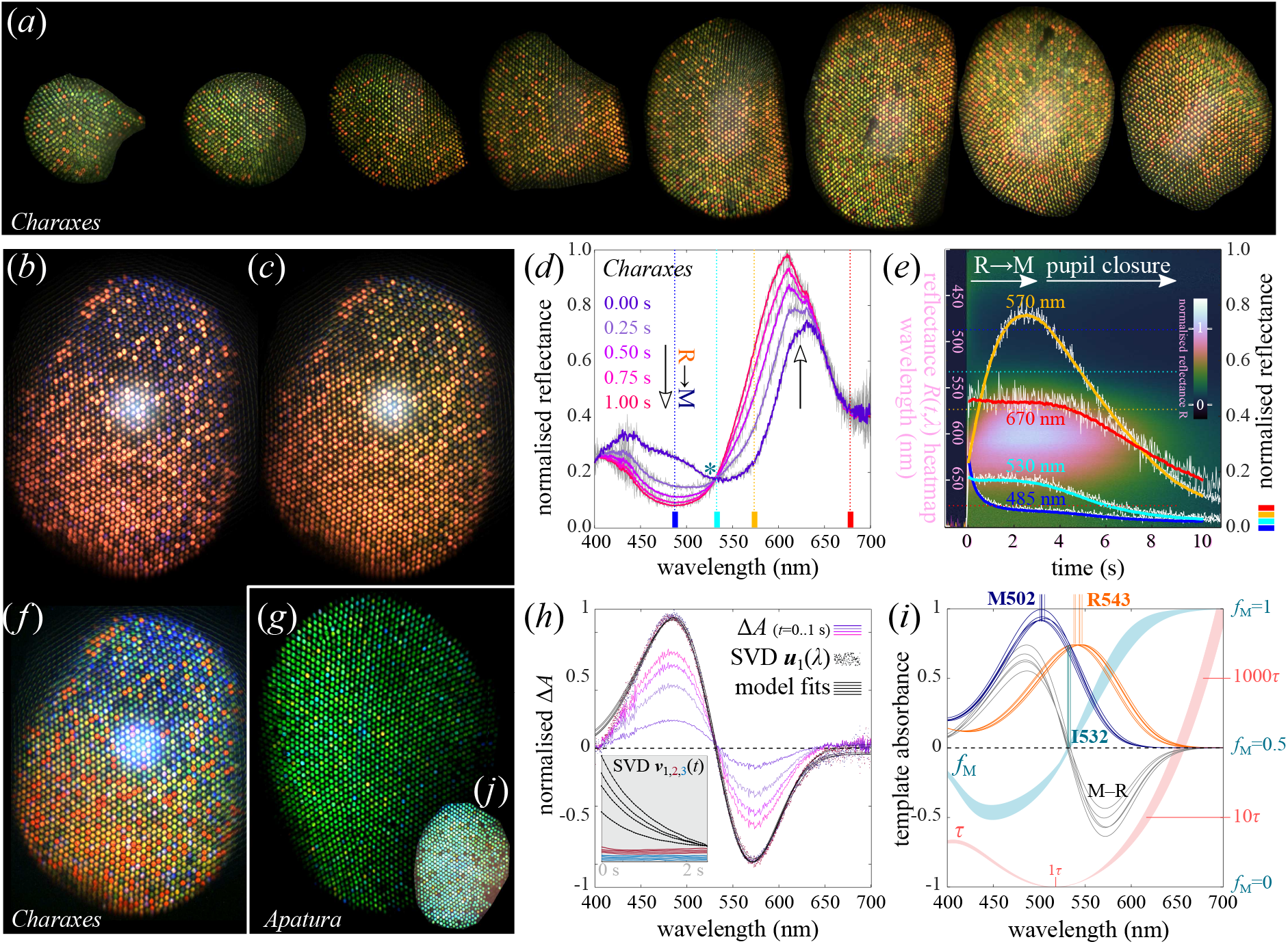
Eyeshine and spectroscopic determination of the LW opsin. **(*a*)** Images of the eyeshine in *Charaxes* with the eye rotated from dorsal to ventral in 15° angles (*left to right*). **(*b***,***c***,***f*)** The eyeshine of *Charaxes*, white-balanced images of (*b*) dark-adapted and (*c*) light-adapted state, (*f*) hyperspectral image of the dark-adapted state. **(*g***,***j*)** The eyeshine of *Apatura*, (*g*) hyperspectral image; (*j*) white-balanced image. **(*d***,***e***,***h***,***i*)** Spectroscopic determination of the main rhodopsin in *Charaxes*. (*d*) Reflectance spectra measured 0.25 s apart (5 traces, *violet-magenta*); raw data as *grey traces*; isosbestic point, *asterisk;* (*e*) Filtered time traces of the eyeshine reflectance at 485, 530, 570 and 670 nm (*blue, cyan, orange, red traces*); raw data (*white* traces); normalised reflectance heatmap *R*(*t,λ*) in the background (y-axis, 400:700 nm); (*h*) Normalised absorbance difference Δ*A* between the dark-adapted spectrum (*t* = 0 s) and the spectra measured at *t* = [0.25, 0.50, 0.75, 1.00 s] (*violet-magenta traces*) from one experiment; SVD spectral components ***u***_1_(*λ*) from five experiments on the same animal (*dots*); fitted template absorbance difference models (*black traces*). Temporal SVD components ***v***_1-3_, ***v***_1_ exhibit an exponential decay with a similar time constant in four experiments (*inset, black traces*). (*i*) fitted absorbance difference (M–R) spectra (*black traces*) from five experiments on one animal; estimated LW rhodopsin (*orange traces*) and metarhodopsin template absorbance spectra (*blue traces*); metarhodopsin fraction *f*_M_ (range 0-1, *cyan shade*), normalised isomerisation time constant *τ*(*λ*) shown with a log y-axis (*red shade*).

When the eye of *Charaxes* was left in the dark for >10 minutes, the dorsal non-red ommatidia were blueish, turning green after a few seconds of illumination (figure 1*b,c* *below*) before the pupil activation would extinguish the eyeshine. The reflectance change had two phases: the first phase was due to rhodopsin-metarhodopsin (R-M) photoisomerisation [31]. Photoisomerisation caused a pronounced reduction of reflectance below the isosbestic point at ∼530 nm (figure 1*d* *below*). The reflectance increase in the wavelength range [550…650 nm] is consistent with the observed eyeshine colour change (figure 1*b,c* *below*). The second phase, due to the pupil closure [32], started about 3 seconds after the light onset and completed about 15 seconds afterwards (figure 1*e* *below*).

The measured reflectance spectra were low-pass filtered, background-corrected and log-transformed. The difference between the first and any later log_10_(*R*) spectrum gave a series of absorbance difference spectra Δ*A* with a peak-trough shape (figure 1*h* *below*). The first 50 log_10_(*R*) spectra were processed using singular value decomposition of the [wavelength×time] matrix. The principal temporal component ***v***_1_(*t*) was decaying exponentially, as expected for a photoisomerisation process. A fit of a template absorbance difference model to the principal spectral component ***u***_1_(*λ*) yielded the estimate for *Charaxes* LW rhodopsin and metarhodopsin, peaking at 543 nm and 502 nm, respectively (R543/M502). The estimated R&M templates, the steady state metarhodopsin fraction *f*_M_(*λ*) and the normalised isomerisation time constant *τ*(*λ*) for monochromatic illumination are shown in figure 1*i* *below*. The estimated LW opsin of *Apatura* is R527/M496 (figure S1 below).

### (b) Intracellular recordings

Photoreceptor identities were revealed by intracellular recordings, using rapid spectral sequence stimulation provided by a fast narrow-band LED source [33]. In both species, we found receptors that depolarised maximally upon ultraviolet or blue stimulation, respectively, and hyperpolarised upon green stimulation (*Charaxes*: figure 2*a,b* *below*; *Apatura*: figure S2 below). We termed the two classes U+G− and B+G−; the letters are signifying the human colours of the unit spectral maxima, {UBGYR} for ultraviolet, blue, green, yellow and red, respectively. In *Charaxes*, we additionally found B+Y− photoreceptors and G+R− photoreceptors (figure 2*c,d* *below*). The responses of the depolarising units B+ and G+ could be isolated by saturating the hyperpolarising units G− and Y− using appropriate long-wavelength adapting illumination (green or orange triangle, middle trace, figure 2*a-c* *below*). Isolation of the hyperpolarising units was less successful due to the sensitivity overlap with the depolarising units in the SW range (purple or blue triangle, bottom trace, figure 2*a-c* *below*). Both units of the G+R− opponent pair could be isolated by selective adaptation of the opponent unit (figure 2*d* *below*), confirming that the sensitivity overlap between the G+ and R− units is minimal [7]. Hyperpolarisation could be enhanced, suppressed or reversed by current injection (figure 2*f* *below*). Both depolarising and hyperpolarising responses were graded along the ∼3 log stimulus intensity range. The aperture of the light stimulus had a minor effect on the hyperpolarisation in the green (figure *S2* *below*). The most frequently encountered cells belonged to the broad-sensitive LW photoreceptor class G and were without a detectable inhibitory input at the retinal level. In these cells, monochromatic adaptation light (of any wavelength) caused a wavelength-independent suppression of the light response (520 nm or 620 nm adapting light used in green and red traces in figure 2*e* *below*).

**Figure 2.**
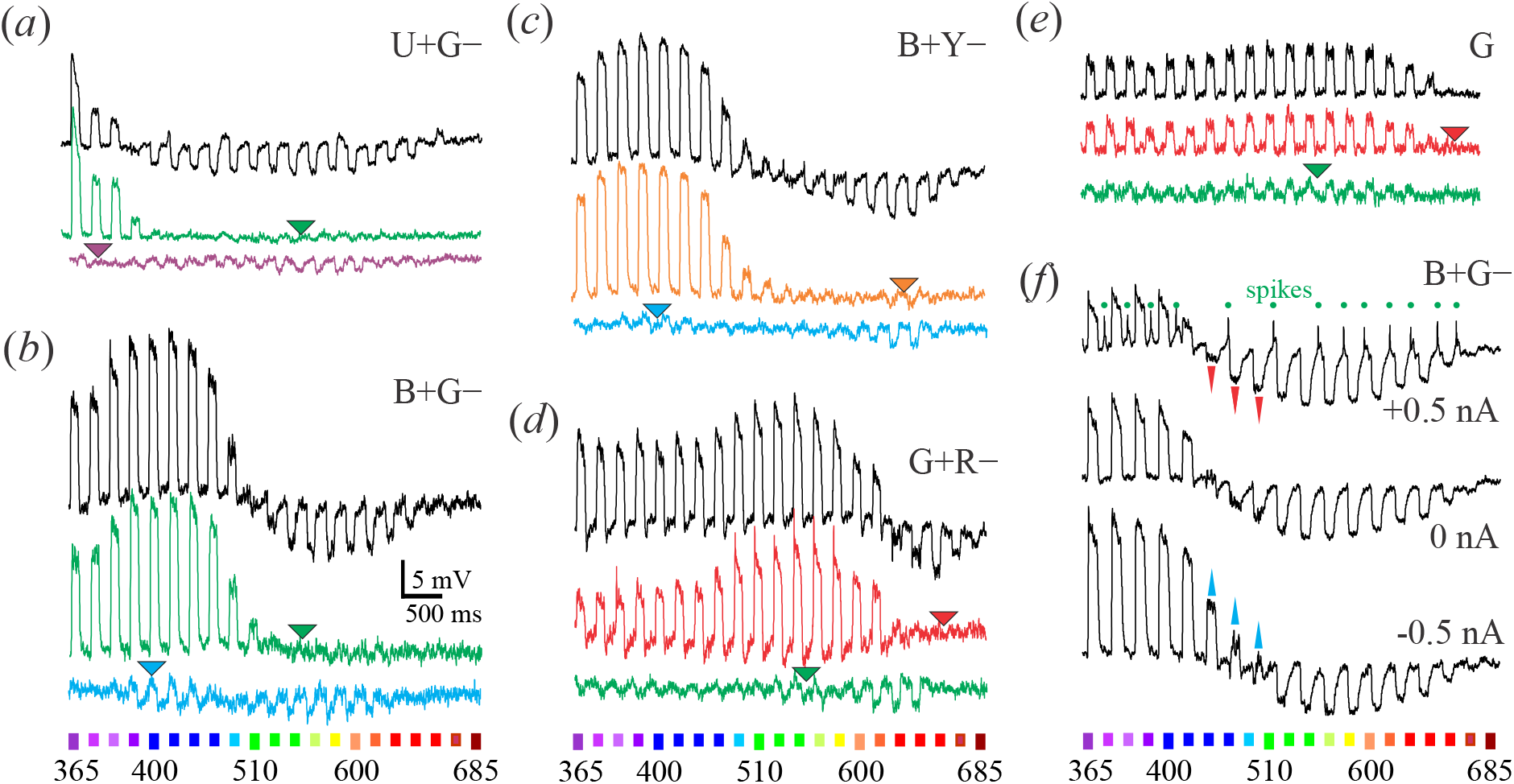
Receptor potentials of *Charaxes* photoreceptors, stimulated with spectral sweeps. **(*a-e*)** Responses in dark-adapted state (black traces) and selectively adapted with steady monochrome light at wavelengths indicated with triangles (coloured traces); **(*a-d*)** opponent cells, **(*e*)** non-opponent cell. **(*f*)** Responses of a B+G– cell at resting membrane potential (0 nA), depolarized (+0.5 nA) and hyperpolarised (−0.5 nA); red and blue arrows indicate reversal of opponent responses. Abbreviations in each panel indicate the cellular spectral class; colour bars at the bottom indicate stimulus (wavelength in nm). Adapting wavelengths: (*a*) 525 and 375 nm, (*b*) 525 and 400 nm, (*c*) 600 and 400 nm, (*d,e*) 625 and 525 nm.

The ancestral set of insect spectral photoreceptors {U, B, G} was found both in *Apatura* and *Charaxes* (figure 3*a,b* *below*; figure S2 *below*). The U and B receptors were maximally sensitive to vertically (parallel to the dorso-ventral body axis) polarised light, consistent with allocation to photoreceptors R1&2. The cells of the main LW receptor class G were maximally sensitive either to horizontally or diagonally polarised light, consistent with the allocation to R3-4 and R5-8, respectively (figure 3*d,e* *below*). In both species, the measured spectral sensitivity of G cells was broader and red-shifted with respect to the corresponding opsin templates with peak sensitivity parameters (*λ*_max_) determined spectrophotometrically (*Apatura*: *λ*_max_ = 527→534 nm; *Charaxes*: *λ*_max_ 543 *→* 548 nm), probably due to self-screening in long photoreceptors [7,15,34]. In *Charaxes*, the additional, yellow-peaking LW receptor class Y (figure 3c below) was similarly consistent with allocation to R3-8 (figure 3*f* *below*). The G+R− receptors, allocated to R1&2, are maximally sensitive to green light and receive opponent input from the red-sensitive, basal R9 that could not be directly impaled [7].

**Figure 3.**
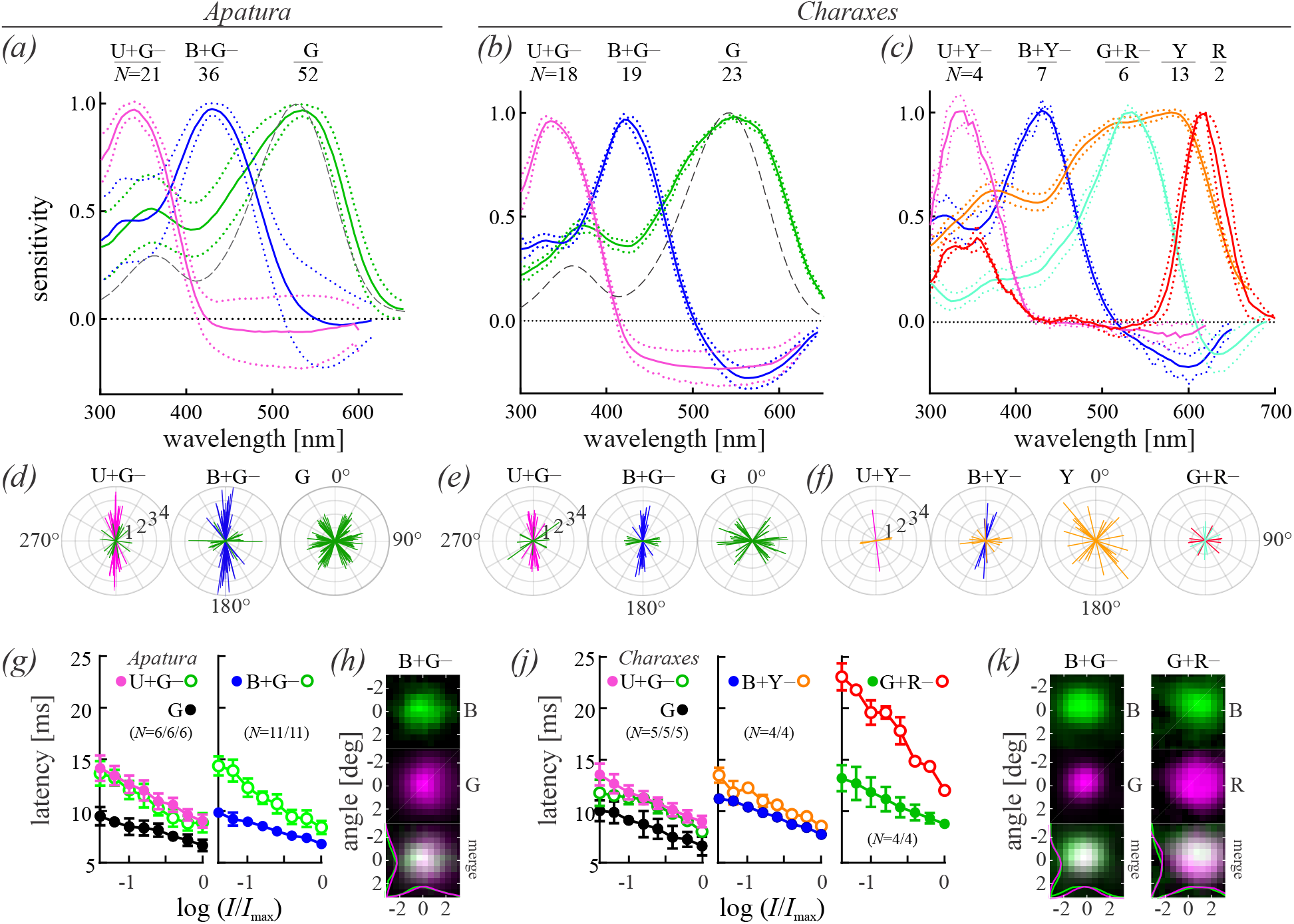
Electrophysiological analysis of photoreceptors in *Apatura* and *Charaxes*. **(*a*-*c*)** Spectral sensitivity (mean±SEM; photoreceptor class and number of analysed cells indicated at the top; dashed grey curves, opsin templates with λ_max_= (*a*) 527 nm and (*b*) 543 nm) in **(*a*)** *Apatura*; **(*b*)** *Charaxes* – basic set, **(*c*)** *Charaxes* – expanded set of photoreceptors. **(*d*-*f*)** Polarization sensitivity (bars indicate magnitude and angle in single cells; polarization sensitivity of opponent units indicated by green bars in U+G–, B+G–, orange bars in U+Y– and B+Y–, red bars in G+R–). **(*g***,***j*)** Response latency (mean±SD, number of analysed cells in brackets) of main cells (full circles), opponent units (empty circles) and non-opponent G cells (black circles) as a function of graded stimulus intensity. **(*h, k*)** Receptive fields of single main cells (top), their opponent units (middle) and overlap between main and opponent units (bottom). (λ_max_ and PS of all spectral classes given in Table 1 below)

The B cells with vertical microvilli were maximally hyperpolarised by their opponent units either around ∼550 or ∼600 nm. The latter is best explained with the opponency of Y units to B cells, yielding the class B+Y− additional to the class B+G−. A similar, albeit more modest distinction could also be made between U+G− and U+Y− classes (figure 3*c* *below*, figure S3 *below*). The receptive fields of the main and the opponent units always overlapped, indicating that the opponency at the level of the retina is not involving pooling from neighbouring ommatidia (figure 3*h,k* *below*).

**Table 1.**
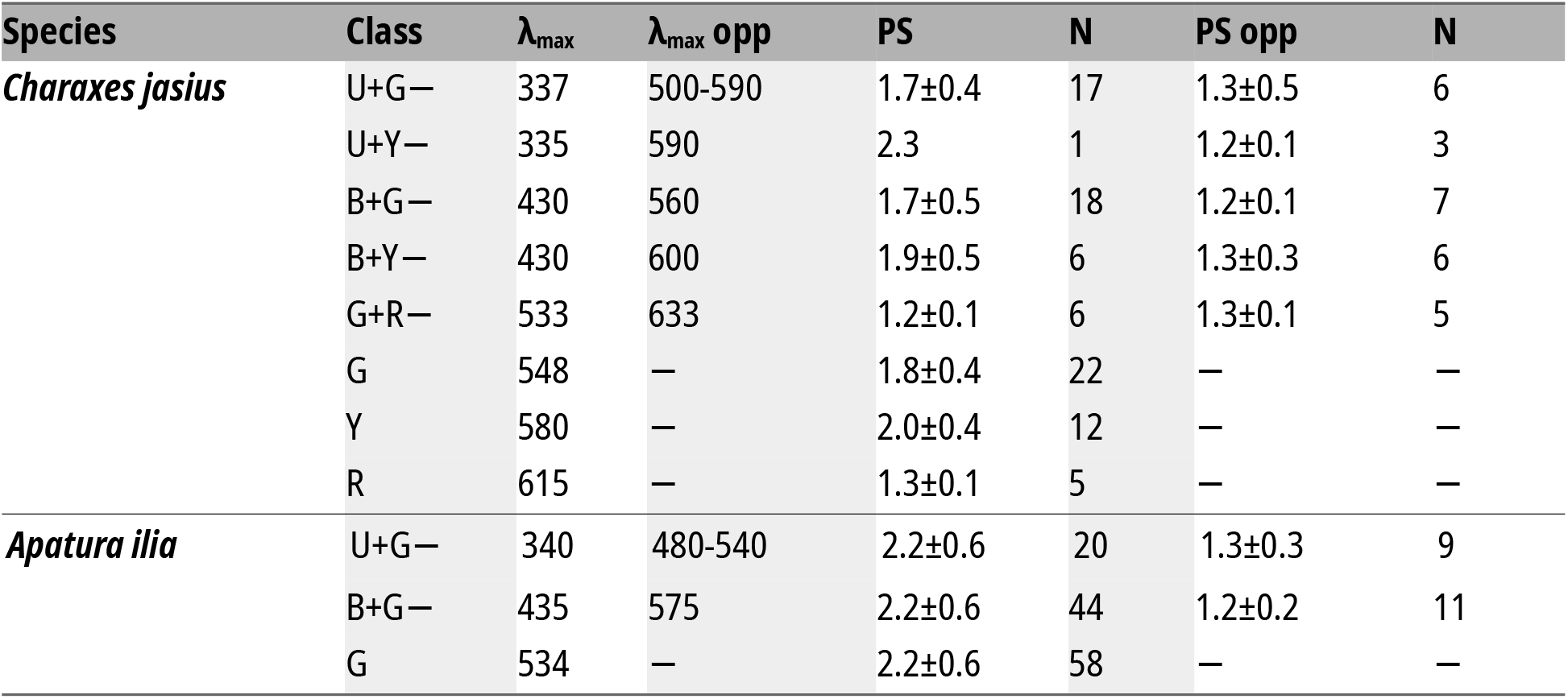
Spectral sensitivity maxima, polarisation sensitivity and angular maxima

Allocation of the main and the opponent units to the receptor positions was further studied by measuring the polarization sensitivity (PS) of the hyperpolarising responses. In most UV+G− and some B+G− cells, PS in the green spectral range was low, suggesting that several green-sensitive cells converge onto a single UV or B cell, so their PS cancels out (figure 3*d* *below*; figure S3 *below*). In a subclass of B+G− and B+Y− cells, the opponent PS was modest (2∼3) and had a horizontal angular maximum, while the main PS was modest to high (2∼4): these opponent cells are good candidates for being the retinal substrate for polarisation vision (figure 3*e* *below*, figure S3 below).

The onset of depolarizing responses was lagging the stimulus onset for 6∼9 ms at the highest stimulus intensity, and for 10∼15 ms when being stimulated with light attenuated by 1.5 log (figure 3*g,j* *above*). The shortest delay was, as expected, in the class G SVF photoreceptors and class Y photoreceptors (*not shown*); the fastest LVF was the class B. Response latency of hyperpolarising units is a sum of the phototransduction and synaptic latency. In B+Y− cells, where the sensitivities of the main and opponent unit are about equal (figure S4 below), the opponent response was delayed relatively to the depolarizing response for ∼1 ms, which is our estimate for the synaptic latency. This latency is consistent with the situation in U+G− class, where the latency of hyperpolarising units was ∼1 ms longer than that of SVF G cells (figure 3*g,j* *above*). In this class, the opponent response was even slightly faster than the depolarising response, possibly due to the slower phototransduction of the U+ unit. The most striking response delay difference was found in G+R− cells, where the opponent response was lagging the depolarising response by 4∼10 ms, likely due to the slow transduction in the minute, light-starved R9 cells (figure 3*j* *above*; figure S5 below).

### (c) Optical retinography

The electrophysiological results in *Apatura* suggested a simple retinal mosaic with two types of LVFs, both receiving opponent signals from G receptors. Physiological evidence suggests that the classes U+G− and B+G− are possibly allocated in pairs to form three types of ommatidia, {UU, UB, BB}, similarly as in *Vanessa* [15]. In the red-eyed *Charaxes*, however, the expanded retinal mosaic contains an additional distal LVF class G+R−, which could, in combination with U and B classes, form three additional ommatidial types, {GG, GU, GB}. The proposed allocation nevertheless awaits molecular validation in both species.

We checked the proposed allocation of six ommatidial types into the eye mosaic of *Charaxes* with optical retinography (ORG) [15], an optical method which reports the compound pupillary sensitivity of the photoreceptors in each ommatidium. We expected that the spectral sensitivity of the pupillary responses, evoked with isoquantal pulses, would resemble the weighted sum of opsin templates in LVFs and SVFs, with relative transduction gains as weights (the gain in cells {U,B} is about tenfold that of cells G; figure S4 below). We note that the pupil is located distally in the retina, where the effects of screening and opponent signals are negligible. Additionally, the compound pupillary response should retain some polarisation sensitivity in the non-red ommatidia in the UV-blue wavelength range, while the polarisation sensitivity of red ommatidia that contain vertical LVF G photoreceptors, would be very small (Table 1 above).

We measured the compound spectral and polarisation sensitivity (PS) of the individual ommatidial pupils in four male specimens of *Charaxes*. Here we present the measurements from 666 ommatidia in the central eye region of a single eye. The pupil sensitivity was measured with an isoquantal spectral sequence (red to UV; UV to red). The sequence was repeated at full, half and quarter intensity. At each wavelength, we acquired bouts of 30 images: the first image was of the dark-adapted eye shine, the remaining were taken after 15 second adapting stimuli of linearly polarised light were delivered to the eye, causing a partial closure of the pupil. The polariser was rotated for 37.25° between stimulations, completing three revolutions in 29 steps. For each bout and each ommatidium, the constant (DC) and modulation (PS) parameters of the pupillary response were estimated using a linear model. The two parameters were normalised to the pupil working range, determined from the images taken in the dark-adapted state and in the fully light-adapted state. The DC and PS spectral sensitivities {*b*_DC_(*λ*), *b*_PS_(*λ*)} were analysed in two passes of singular value decomposition and then classified using k-means clustering (see Methods). The six ommatidial clusters that formed (figure 4*i* *below*) had compound pupillary spectral and polarisation sensitivities consistent with the three basic ommatidial types {BB, UB, UU} and with the proposed ommatidial types containing green-sensitive LVFs {GB, GG, GU}.

**Figure 4.**
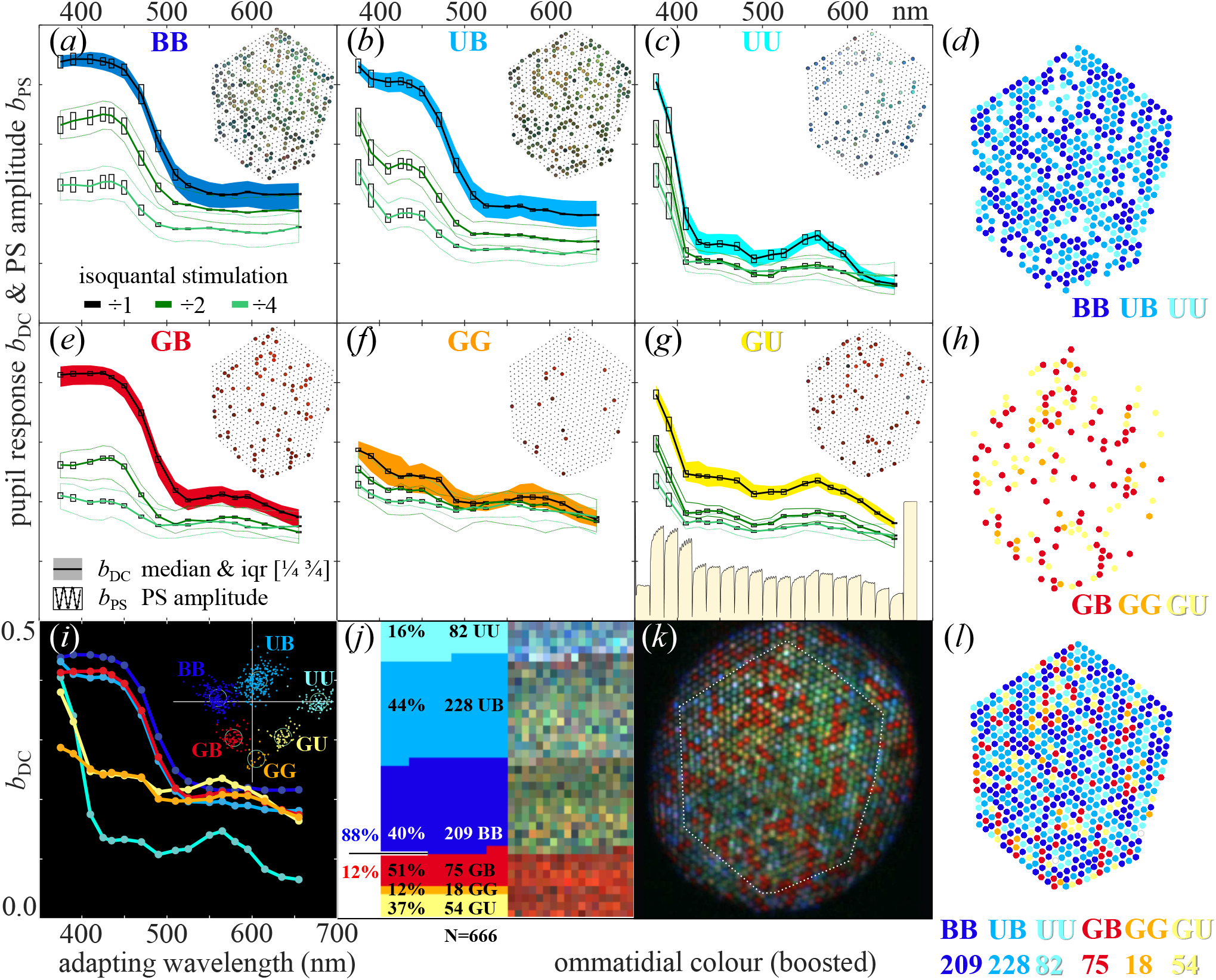
Optical retinography of *Charaxes jasius*. **(*a***,***b***,***c***,***e***,***f***,***g*)** Pupil sensitivity spectra of the six ommatidial clusters, BB (*a*), UB (*b*), UU (*c*), GB (*e*), GG (*f*) and GU (*g*), obtained with isoquantal adapting light (unattenuated: *black*, half intensity: *dark green*, quarter intensity: *light green*). The pupil responses, expressed as reduction of ommatidial reflectance (0 = no response, 1 = maximal reduction obtained with a saturating broadband stimulus) are shown as cluster’s median (*solid lines*) and inter-quartile range (*shaded area*). Median pupil polarisation sensitivity amplitudes (modulation of the response obtained at different orientations of the linearly polarised adapting light) are shown with *boxes*. The pupil response y-axis range is [0.0 … 0.5]. Allocation of the types to the ommatidial lattice is shown in *insets. (****d***,***h***,***l***) Allocation of the ommatidial types {BB,UB,UU} (*d*), types {GB,GG,GU} (*h*), and all six types (*l*) to the ommatidial lattice, cluster member counts at the bottom. (***i***) plot of the pupil sensitivities of the six ommatidial clusters to unattenuated adapting stimuli; scatter plot of the principal two clustering scores of 666 ommatidia (*inset*). (***j***) fractions and counts of ommatidia in clusters (*left*), boosted ommatidial colours (*right*) obtained from the hyperspectral image. (***k***) hyperspectral image of the dark-adapted ommatidial lattice.

The constant (DC) pupil responses of the ommatidial clusters *b*_DC_(*λ*) are shown as median and inter-quartile range, the median of PS modulation *b*_PS_(*λ*) is depicted with bars (figure 4*a-c, e-g* *below*). The clusters {BB, UB} had pronounced PS in the UV-blue spectral part figure 4*a,b* *below*). The cluster UU had a high PS across the whole spectrum (figure 4*c* *below*). The remaining three clusters had generally lower PS (figure 4*e-g* *below*). The distribution of the six clusters in the ommatidial lattice seems to be random (figure 4*d,h* *below*). The boosted colours measured from the hyperspectral image of the eyeshine (figure 4*k* *below*) were mapped to the six clusters. The two most numerous clusters {UB,BB} were green-shining, the cluster {UU} was blueish (figure 1*b* *above*, 4*j below*). Most notably, the three least numerous clusters with low PS (figure 4*e-g* *below*) were all red-shining (figure 4j below). The two more numerous red clusters are consistent with ommatidial types {GB, GU}, the least numerous cluster is likely of type GG (figure 5 below).

**Figure 5.**
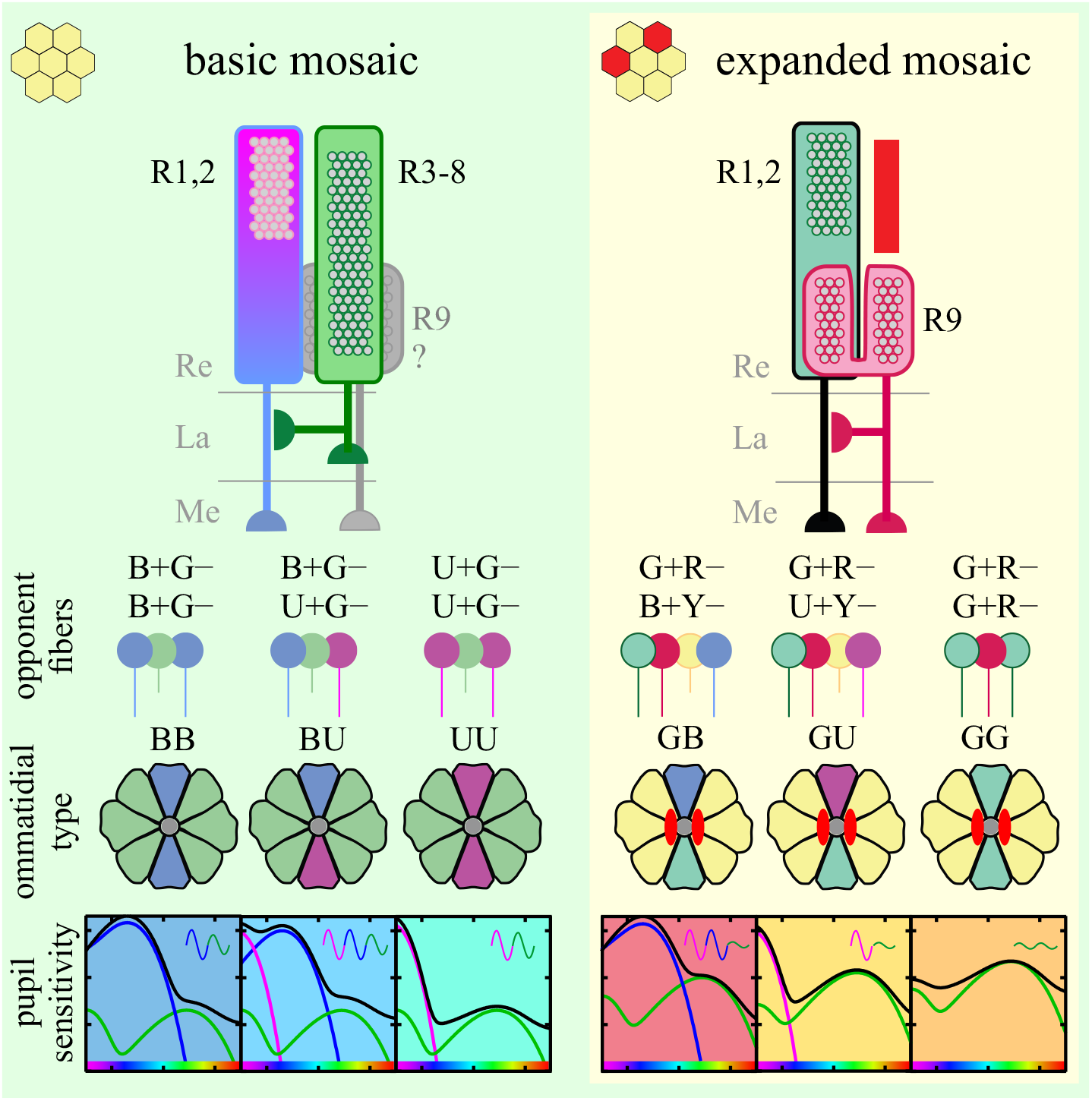
Proposed allocation of opponent photoreceptors in the retinal mosaic. **Top**, cellular identity of LVF (R1, 2, 9) and SVF (R3-8) photoreceptors in opponent pairs; middle, their allocation into ommatidial types (not yet confirmed with opsin mRNA or protein localisation). U, B, G, Y, G+R – and R receptors are coloured pink, blue, green, yellow, teal and red; microvilli and rhabdoms are depicted with grey circles; red stripe above R9 and red ovals next to the rhabdoms indicate the red screening pigment. **Bottom**, spectral sensitivity of pupil in each ommatidial type, a weighted sum of U, B and G LVF spectral sensitivities, superimposed on a background response of SVFs; the contribution of LVFs is scaled according to the transduction gain; the background is coloured as the ommatidial types in Fig. 4*a-g*. Sinusoidal insets indicate polarization sensitivity of the pupil in UV, blue and green (pink, blue and green curves); spectral range indicated with rainbows on the x-axis). Re, retina; La, lamina; Me, medulla.

## III. Discussion

We have shown that in *Apatura ilia*, a nymphalid butterfly with a basic set of spectral photoreceptors {U, B, G}, the retina is likely built from three ommatidial types without red pigments that form a simple mosaic. In *Charaxes jasius*, a nymphalid with an expanded set of LVF photoreceptor classes {G+R−, R} and a SVF photoreceptor subclass Y, the retinal mosaic is complex, having three additional ommatidial types, allocated to the red ommatidia (figure 5 below).

We found similar expanded spectral sets of photoreceptors in other species with red ommatidia and with a complex mosaic eye shine, e.g. the monarch (*Danaus plexippus*), blue morpho (*Morpho peleides*) and prepona (*Archaeoprepona demophon*) (figure S6 *below*). Interestingly, these butterflies likely have only three opsin genes {U, B, G=LW} [35,36], suggesting that the expanded set of receptors {G+R−, Y and R} is implemented on the basis of optical filtering of a single LW opsin. The simple retinal mosaic can support trichromatic vision in the ultraviolet to green range [28,37]. Colour discrimination in the red wavelength range and the putative tetrachromatic vision are only supported in the eyes with a complex retinal mosaic, confirmed behaviourally in *Danaus* or *Heliconius* [28,38], but not yet in *Charaxes*. The retinal complexity is likely costly and seems to be evolutionarily switched on or off, depending on the visual ecology of the species. Extension of colour vision into the red wavelength range in brush-footed butterflies is associated with the simultaneous occurrence of multiple features of the visual system: the green-sensitive R1&2, the red screening pigment, and a functional R9 with direct retinal opponency to the class G+R− LVF. These features are likely controlled by an additional stochastic genetic switch similar to the known *spineless* transcription factor [16].

Quite possibly, the expanded retinal mosaic is also associated with a red-shifted LW opsin. Nymphalid LW opsin is subjected to extensive evolutionary tuning, with λ_max_ varying between 515 nm and 565 nm [35], but the physiological relevance of this tuning is unknown. In nymphalids with a uniform eyeshine, the LW opsin tends to peak below 530 nm [27], whereas in the red-eyed butterflies, the LW opsin’s peak tends to shift above 540 nm. A similar shift has been implicated in the evolution of red colour vision in lycaenid butterflies [39].

We note that a basal photoreceptor, receiving light filtered by red screening pigments can have high sensitivity and a high signal-to-noise ratio only by expressing a red-shifted LW opsin. Assuming narrowband red light (620 nm), R9 would be approximately twice and five times more sensitive, if R525 is red-shifted for 10 and 25 nm, respectively. Photoconversion of metarhodopsin seems to be very ineffective in a red-sensitive R9 (figure 1*i* *above*), so the cell probably relies on enzymatic pigment conversion to maintain high sensitivity.

The simple nymphalid retina contains two colour-opponent channels (here U+G− and B+G−) and an achromatic channel (here SVF, G), following the design principles for optimal information transfer in trichromatic vision [3]. In the expanded retina, a new opponent channel (G+R−) and a Y subclass achromatic channel are added, likely on the basis of the same LW opsin as the green-sensitive R3-8. The channel expansion is implemented through a red screening pigment that tunes the basal red receptor R9. Three interesting functional features can be elucidated. (1) LVF R1,2 can receive opponent inputs from either LVF or SVF (U&B from R3-8, G from R9). (2) Opponency seems to be unidirectional, LVFs are not opponent to SVF. (3) Opponent cells (G SVFs and R9 LVF) have a red-shifted sensitivity, compared to their postsynaptic partners R1&2. (We note that due to the absence of direct recordings, we cannot exclude the possibility that R9 are receiving an opponent input from another class of SVF.) This implementation is quite different from *Papilio*, where direct opponency is present among all kinds of visual fibres, including LVFs being opponent to both LVFs and SVFs, and SW-peaking classes being opponent to LW classes [5], or from *Drosophila*, where direct opponency is only present between the LVFs [4].

The SW photoreceptors U and B have a higher phototransduction gain (effectively more millivolts of depolarization per absorbed quanta) than the LW photoreceptors (figure S4 *below*). Direct opponency from U&B LVF would cause strong inhibition of G&Y SVF. This would be detrimental for the signal-to-noise ratio in the achromatic visual pathway relayed *via* the lamina monopolar cells (LMCs). The opponent signalling from SW to LW is likely implemented at a later stage in the visual pathway, probably *via* interneurons in the medulla [8,9]. In the complex nymphalid retina with red ommatidia, the tiny, light-starved, high-gain R9 cells can utilise a novel, ***private opponent channel*** in the form of green-sensitive LVF R1&2, thereby avoiding to send an opponent signal into the R3-8 achromatic pathway.

The LVFs in our study exhibited various levels of hyperpolarisation due to the opponent cells; generally, the opponency in *Apatura* was much less pronounced than in *Charaxes* (figure 2*a-c* *above*). The sensitivity of the receptors in opponent pairs must be tuned so that the degree of inhibition is *just right*. The inhibition should not be too weak, or else spectral discrimination would not work, nor too strong, to the detriment of signal generation and propagation. We hypothesize that the inhibition gain may be adjustable by yet unknown mechanisms that depend on light adaptation and internal state-dependent efferent inputs from the central nervous system. Interestingly, many LVFs from all spectral classes exhibited slow spikes (figure 2*f* *above*, figure S2*a* *below*), that are very loosely correlated with light stimulation and depolarization. These spikes may have been initiated by the adjacent cells that may be modulating the physiological properties of their impaled neighbour. Further comparative research is needed to assess how do the opponent pairs respond to the changes in spectral composition of ambient light across different time scales, and what are the optimal spectral opponent combinations for the particular photic environment.

## IV. Experimental methods

### (d) Animals

The butterflies were collected near Zadar and Mali Lošinj, Croatia (*C. jasius*) or bred and shipped from the UK (*A. ilia*) by Mr. Mark Youles as a part of ongoing collaboration. Adult butterflies were kept at 27°C and 80 % relative humidity and regularly fed sucrose solution.

### (e) Rhodopsin isomerisation spectroscopy

The template spectra of the LW rhodopsin and metarhodopsin can be estimated using a spectroscopic method employing a broadband light source both for isomerisation and the measurement [40]. The animal was immobilised in a pipette tip attached to a manual goniometer and placed under a Leitz Orthoplan microscope with a custom epi-illumination attachment with a 50 % beamsplitter. The light source was a modified violet-chip based white LED (Soraa MR16-50-B03) driven with a constant current source. A long working distance objective (either Nikon Plan ELWD 20xNA0.40 1.2/160, part 120152, or an Olympus MPlan 10xNA0.25 0/∞) was focussed on the eye’s curvature centre. A Bertrand eyepiece was used to finely position the animal and set the illuminator’s aperture and field iris so that the whole objective back-focal plane was filled with ommatidial reflections. The eye’s luminous deep pseudopupil was imaged onto the iris plane of a custom-made microspectrometer head and relayed to a rosette-to-line fibre bundle attached to a spectrometer (Ocean Optics USB2000) controlled from GNU Octave. Eyeshine images were taken with a Raspberry Pi4, equipped with a Pi HQ Camera (RGB CMOS, 20MP).

Before each measurement, the animal was left to dark adapt for 10-60 minutes, so that the visual pigment would be predominantly in the rhodopsin isoform. The shutter was opened for 10∼15 seconds, and several hundred spectra with integration time 15∼25 ms were taken. Electronic (thermal) noise and the animal or objective background were subtracted from the spectral series. Data processing was implemented in GNU Octave with Signal and Optim toolboxes, using singular value decomposition, where a matrix with a temporal series of spectra **M** is decomposed into *n* spectral (**U**) and temporal (**V**) components, **M**_[λ×t]_→ **U**_[λ×n]_**D**_[n×n]_**V**_[t×n]_^T^.

The raw data matrix **M**_0_ with count spectra [2048 pixels × 500 acquisitions] was decomposed into spectral and temporal components that were separately low-pass filtered using a zero-phase filters (function *filtfilt*). A filtered matrix **M**_1_ was recomposed from the first few signal-bearing components, and then log-transformed, **M**_2_ = log_10_(**M**_1_). The first 50 spectra of **M**_2_ were again decomposed. The fundamental component {***u***_0_,***v***_0_} was approximately constant in time and could be discarded, while the next temporal component vector ***v***_1_ followed an exponential relaxation course, indicative of a photo-isomerisation process. The corresponding spectral component vector ***u***_1_, analogous to the absorbance-difference spectrum was fitted with a template absorbance difference model based on Govardovskii [41] templates ***Γ***_*λ*_, ***û***_1_ = *a* (*c* ***Γ***_*M*_ *−* ***Γ***_*R*_) + (*b / a*), where *c* is M/R peak absorbance ratio, *b* is background correction, and *a* is a technical scaling parameter. We used nested models where the parameters could be constrained to *b* = 0 or *c* = 1.25, the latter being a biblical value for M/R peak absorbance ratio. A model fit yielded *λ*_R_, *λ*_M_ and optionally *c*. In the case of *Charaxes*, the value *c* = 1.25 seems to be correct, while the experimental data for *Apatura*, due to close-lying R&M peaks, does not allow for a reliable estimation of *c*. The data points below 430 nm were excluded from the fit due to a systematic deviation (see also Figure S1 below).

### (f) Electrophysiological recordings

Intracellular recordings were performed as described previously [7]. Briefly, immobilized animals were placed with the head in the centre of rotation into a goniometer that also carried the micromanipulator with sharp electrodes (Sensapex, Oulu, Finland). The recordings were performed with an amplifier (SEC-10LX, NPI, Tamm, Germany) in bridge mode or discontinuous clamp mode at 20 kHz & 0.25 duty cycle. The electrodes were pulled from borosilicate glass and had resistance in the range 80∼120 MΩ when filled with 3M KCl. Light stimulation was provided by a 75 W XBO lamp (Cairn Research, Kent, UK), filtered through a motorized monochromator (B&M Optik, Limburg, Germany), a computer-controlled neutral density wedge filter (Thorlabs, Dachau, Germany) and, for polarization sensitivity measurements, a UV-capable polariser (OUV2500, Knight Optical, UK). The second source was a ‘LED Synth’ with narrow-band LEDs between 365 and 685 nm in 15 nm intervals. Both sources were combined with a beam splitter (Thorlabs, Dachau, Germany) and isoquantised with a irradiance-calibrated spectrophotometer (Flame, Ocean Optics, USA) to yield equal photon (isoquantal) flux density at all wavelengths (max. 1.5×10^15^ photons cm^−2^ s^−1^). An iris allowed to adjust the aperture of the coaxial stimulating beam between 1.5° and 20°. Receptive fields of the photoreceptors were mapped with an RGB DLP projector (LightCrafter 4500, Texas Instruments, USA) that projected to a back-projection screen (ST-Pro-X, Screen-Tech e.K., Hohenaspe, Germany) at a refresh rate 220 Hz using software package PsychoPy.

### (g) Eyeshine

Eyeshine was observed with a custom epi-illumination microscope built from Thorlabs, Edmund Optics and Linos parts as described elsewhere [15,29]. The relaying lenses were near-UV achromatic doublets (Edmund Optics). The main objective lens was a Zeiss LD-Epiplan 20×NA0.40 objective (part 442840). Images were taken with monochrome or RGB CMOS cameras (1.6, 2.3, 20.0 MP, BFS-U3-16S2, BFLY-U3-23S6, BFS-U3-200S6, all FLIR/PointGray).

The illumination for RGB images was provided by a white LED (colour temperature ∼ 3000 K), filtered by a purplish filter (Lee Filters) that brought the three colour channel gains close to unity. Hyperspectral images were taken with a monochrome camera, using the LED synth [33] was used as the light source. Exposure and gain were optimised for each image. The instrumental background, due to reflections from the objective lenses, was acquired separately, averaged and subtracted from the image series. Images were processed in ImageJ/Fiji [42]. Background correction was performed by subtracting an averaged image of lens reflections without the animal. The stacks were aligned using StackReg [43]. Substacks covering spectral ranges where a similar eyeshine pattern was observed (e.g. 380-440 nm, 480-540 nm, 600-660 nm) were averaged and joined into a pseudo-coloured image (figure 1*f* *above*, figure 4*k* *above*).

### (h) Optical retinography

Optical retinography is an optical method reporting the compound sensitivity of the photoreceptors in each ommatidium [15]. We measured the pupillary action spectra of ommatidia in the central eye region of *Charaxes*. The experiments were performed in the epi-illumination microscope equipped with a monochrome CMOS camera as described above. The light source for test images was a white LED, filtered with an orange or red-coloured glass long-pass filter. The adapting light source was the LED synth [33] with 20 LEDs covering the range 365∼660 nm. The combined beam was depolarized using a liquid crystal depolarizer (DPP25-A, Thorlabs) before being focussed to a 1 mm diameter, NA 0.40 polymer fibre. The fibre output was collimated with an aspheric lens and sent through a rotatable, UV-capable polariser.

The adaptation experiment was conducted three times, at full, half and quarter intensity of the isoquantal adapting light source. In each of the experiments, we acquired m = 1200 images, divided into 40 bouts. In each bout, we recorded a dark-adapted image and 29 images where monochrome, isoquantal, linearly polarised adapting light was switched on for 15 seconds prior to taking the test image. The polariser was rotated for 37.25 ° between the images, so the 30^th^ image completed the third rotation in the same angular position as the 2^nd^ image. Each bout was followed by a one minute dark adaptation period. The bouts #1 and #21 were dark-adapted control, the bouts #20 and #40 were control with saturating adaptation light. The bouts [#2…#19] and [#22…#39] were sequences with spectral adaptation going from red to UV, and from UV to red, respectively. We reversed the bouts [#2…#19], yielding six experimental runs with adaptation wavelength going from UV to red.

The ROI of ommatidia were found using Fiji/ImageJ [42] functions “Find Maxima” and “Analyse Particles”. The adaptation states of n = 666 ommatidia in m = 3600 images (6 experimental runs × 20 bouts × 30 images) were exported as a table of average ROI grey values *g*_[n×m]_. The pupil range of each ommatidium was obtained by finding the maximal grey value in the fully dark-adapted state *g*_n,DA_ and the minimal grey in the fully light-adapted state *g*_n,LA_. The relative pupil activation for the ommatidium *n* in image *m* was calculated as *p*_n,m_ = 1 − (*g*_n,m_−*g*_n,LA_)/(*g*_n,DA_ − *g*_n,LA_).

The 30 pupil responses in each bout, ***p***_n,m_ = [*p*_n,m_ … *p*_n,m+30_]^T^ were fitted with a linear model ***p*** = **A*b***. The matrix **A**_[30×5]_ was constructed from column vectors containing a constant (DC) component, sine and cosine modulation, linear trend and exponential decay with a predetermined time constant. Solving the matrix equation ***b=*A*\p*** yielded a parameter vector containing the constant pupillary response parameter *b*_DC_ and the oscillating parameters {*b*_cos_, *b*_sin_}. From the latter two, PS modulation amplitude was calculated as *b*_PS_ = 2|(*b*_cos_ + î*b*_sin_)|; we did not analyse the PS phase. The parameters *b*_DC_ and *b*_PS_ obtained in each of the six experimental runs were placed into matrices **B**_DC_ and **B**_PS_ of size [666 ommatidia × 18 wavelengths]. Each of these 12 matrices were separately normalised and decomposed using singular value decomposition (SVD), yielding score and spectral matrices. The first three score vectors [666×3] from each decomposition were concatenated, yielding a matrix of size [666 ommatidia × 36 scores], coming from 2 parameters {*b*_DC_, *b*_PS_} × 6 experimental runs × 3 score vectors. This matrix was again decomposed, giving the final score matrix with 5∼6 signal-bearing score vectors that was used for k-means clustering. The first two score components are shown as the scatter in inset of Figure 4*i* *above*.

## V. Acknowledgements

The authors wish to thank the company Votan d.o.o. for the technical development of the LED Synthesisers. The authors are grateful to Mark Youles for providing the pupae of *Apatura ilia*, and to Doekele Stavenga for reading the manuscript. This work is dedicated to the late Guillermo Zaccardi, whose pioneering work on colour vision of nymphalids was a source of inspiration for this study.

This study was financially supported by the AFOSR/EOARD (grant FA9550-15-1-0068, to GB&MI), by Slovenian Research Agency (P3-0333 to GB, AM&PP) and jointly by the European Regional Development Fund and MESS of Slovenia (grant 5442-1/2018/434 to PP).

## Author Contributions

Study design GB&PP; Optical measurements PP; Electrophysiology GB&MI; Anatomy AM; Data analysis GB PP MI; Writing – original draft GB&PP; Writing – review & editing GB PP MI AM. The authors declare no competing interests.

## VI. Supplementary Information

### i. Technicalities of rhodopsin spectroscopy & template fitting

**Figure S1.**
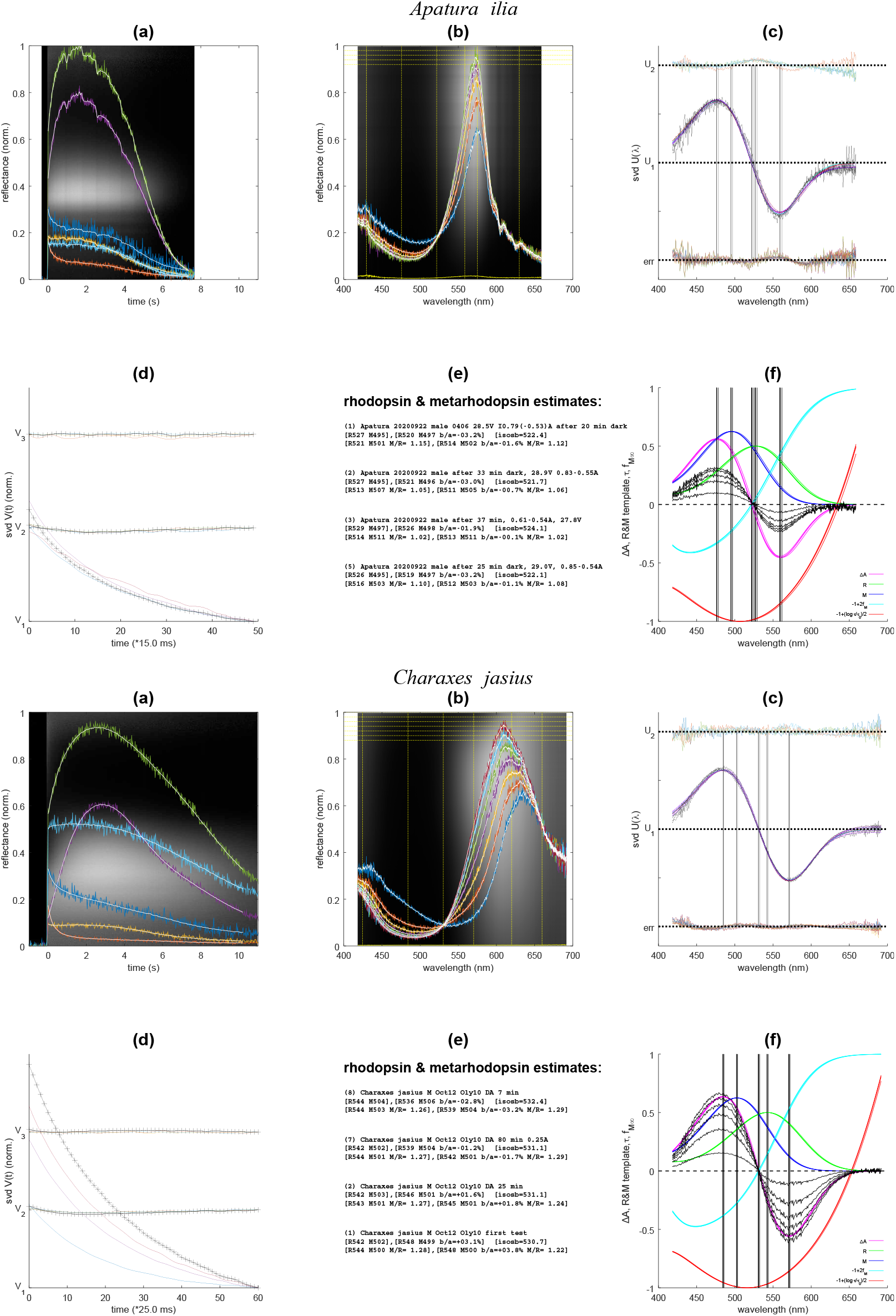
Spectroscopic determination of LW rhodopsin in *Apatura ilia* and *Charaxes jasius*. **(*a*)** Time traces of the eyeshine reflectance change (6 coloured traces, measured at the wavelengths designated in (*b*), the trace corresponding to the isosbestic point is shown in *yellow*); filtered data is shown as white traces; waterfall y-axis [400:700 nm]. **(b)** Reflectance spectra measured 0.25 s apart (*5 coloured traces*); filtered data (*white traces*); instrumental background (yellow trace). **(c)** Components ***u***_1_(*λ*) (*black* Z-shaped traces) obtained from four experiments, sixteen fit curves (4 experiments × 4 models), fit errors (*coloured traces, bottom*), and components ***u***_2_(*λ*) (*coloured traces, top*). Fit errors show a consistent bias below 430 nm, possibly due to using inadequate templates, due to interference from SW opsins, or due to the complications arising from the intricacies of the optical measurement, signal processing, or the eye optics itself. **(d)** Temporal components ***v***(*t*) of SVD absorbance spectra, multiplied by signal energies obtained from the diagonal matrix. Component ***v***_1_ has a course consistent with a first-order exponential relaxation. Component ***v***_2_ exhibits exponential relaxation, faster than ***v***_1_, and is perhaps due to isomerisation of SW rhodopsins; ***v***_3_ is negligible. The signal energy in ***v***_1_ is dominating, and is proportional to the absorbance difference change in a particular experiment, which in turn depends predominantly on the dark adaptation interval used between the experiments. **(e)** Numerical estimates for R, M, baseline shift, and M/R peak ratio obtained from the four nested models. For *Charaxes*, all four models agree well. The free fit for M/R peak ratio confirms the biblical value 1.25. For *Apatura*, only the first two models (with constrained M/R ratio) are consistent. **(f)** Templates obtained from the parameters estimated from the fits of the minimal model to the data. Rhodopsin templates (*green*), metarhodopsin templates (*blue*), absorbance difference (*magenta*), steady state metarhodopsin fraction *f*_M_(*λ*) (i.e. the fraction obtained upon sufficiently long illumination with monochrome light of wavelength *λ*), scaled as *y* = 2*f*_M_−1 (*cyan*); the associated normalised time constant of isomerisation *τ*(*λ*), scaled as *y* = ½(log_10_ *τ*/*τ*_min_) − 1; the *y* values [-1.0, -0.5, 0.0, 0.5, 1.0] correspond to (log *τ*/*τ*_min_)=[1, 10, 10^2^, 10^3^, 10^4^] (*red*).

**Table S1:**
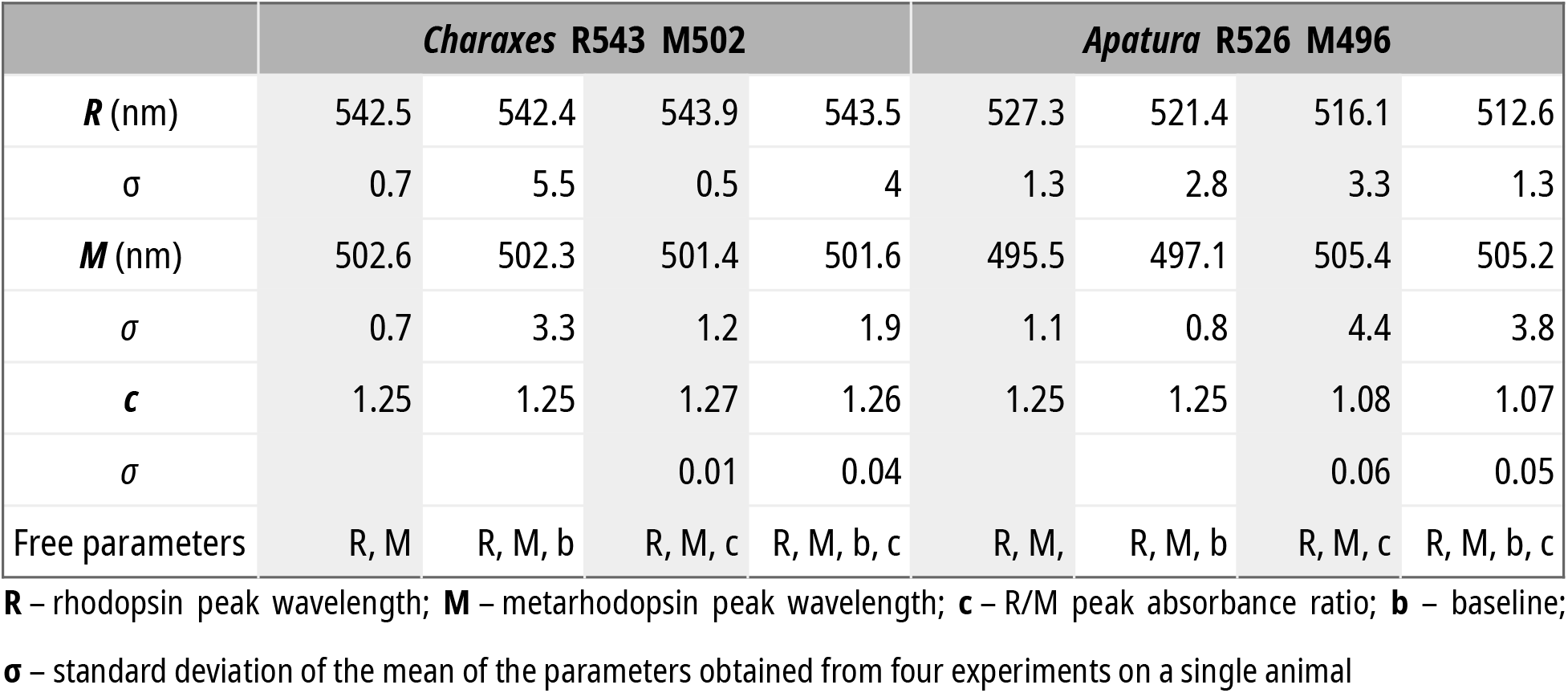
fit parameters

|  | *Charaxes* R543 M502 |  |  |  | *Apatura* R526 M496 |  |  |  |
| --- | --- | --- | --- | --- | --- | --- | --- | --- |
| **R** (nm) | 542.5 | 542.4 | 543.9 | 543.5 | 527.3 | 521.4 | 516.1 | 512.6 |
| $\sigma$ | 0.7 | 5.5 | 0.5 | 4 | 1.3 | 2.8 | 3.3 | 1.3 |
| **M** (nm) | 502.6 | 502.3 | 501.4 | 501.6 | 495.5 | 497.1 | 505.4 | 505.2 |
| $\sigma$ | 0.7 | 3.3 | 1.2 | 1.9 | 1.1 | 0.8 | 4.4 | 3.8 |
| **c** | 1.25 | 1.25 | 1.27 | 1.26 | 1.25 | 1.25 | 1.08 | 1.07 |
| $\sigma$ | | | 0.01 | 0.04 | | | 0.06 | 0.05 |
| Free parameters | R, M | R, M, b | R, M, c | R, M, b, c | R, M, | R, M, b | R, M, c | R, M, b, c |

**R** – rhodopsin peak wavelength; **M** – metarhodopsin peak wavelength; **c** – R/M peak absorbance ratio; **b** – baseline; $\sigma$ – standard deviation of the mean of the parameters obtained from four experiments on a single animal

### ii. Basic opponent classes, stimulation aperture and intensity

**Figure S2.**
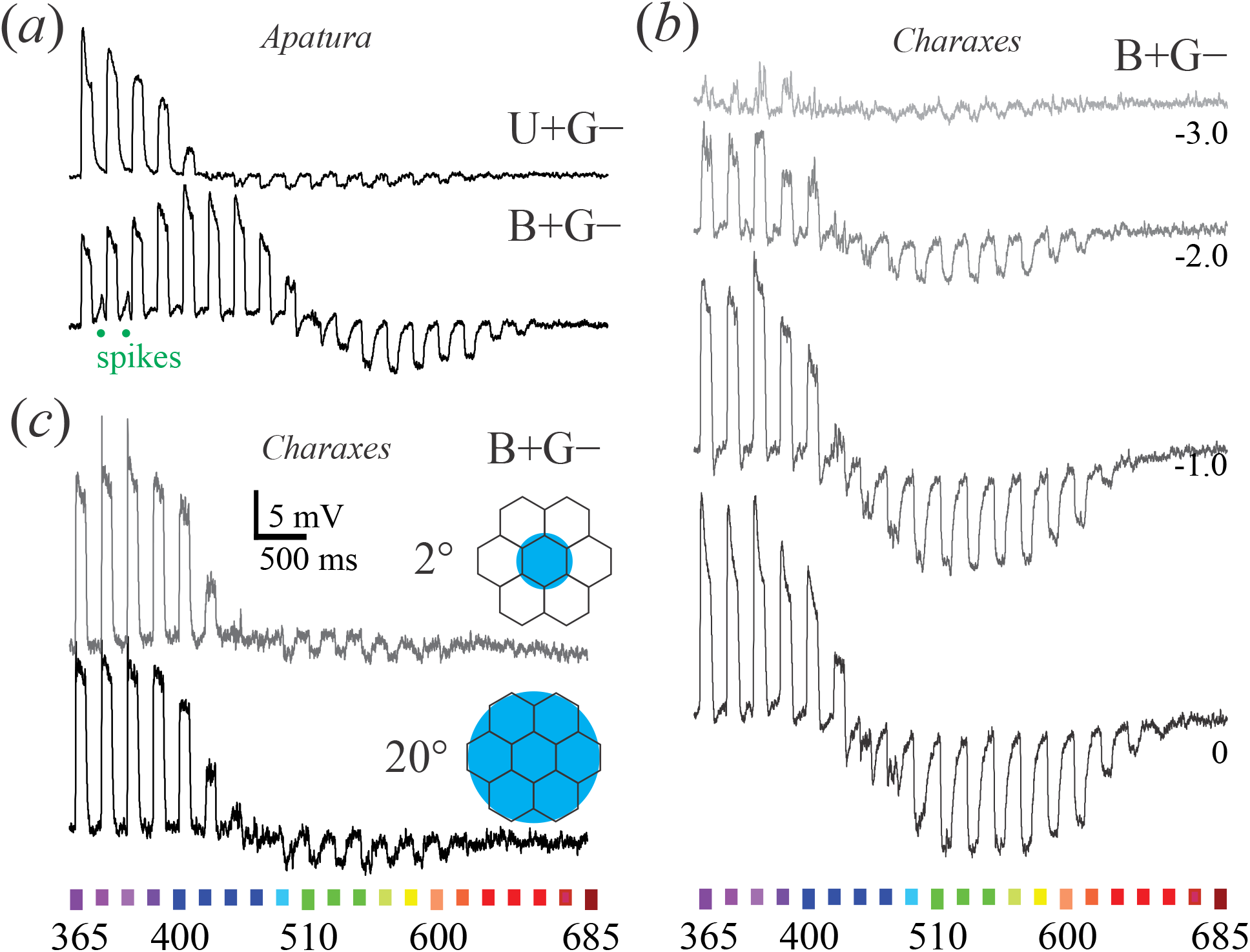
Photoreceptor responses to spectral sweeps of light flashes. ***(a)*** U+G- and B+G- opponent receptors in *Apatura* ***(b)*** Responses of a B+G- opponent receptor in *Charaxes* to sweeps with light intensity graded between – 3 log (10^−3^) and 0 log (full intensity); opponent signals are present already at the lowest intensity. ***(c)*** Responses of a B+G- opponent receptor in *Charaxes* to two sweeps incident into a single ommatidium (2° aperture) and ∼ 50 ommatidia (20° aperture); opponent signals are only slightly affected by the aperture angle of illumination. *Colour bars* depict approximate human colours of stimuli, wavelengths in nm.

### iii. Spectral and polarisation analysis of the opponent pairs

**Figure S3.**
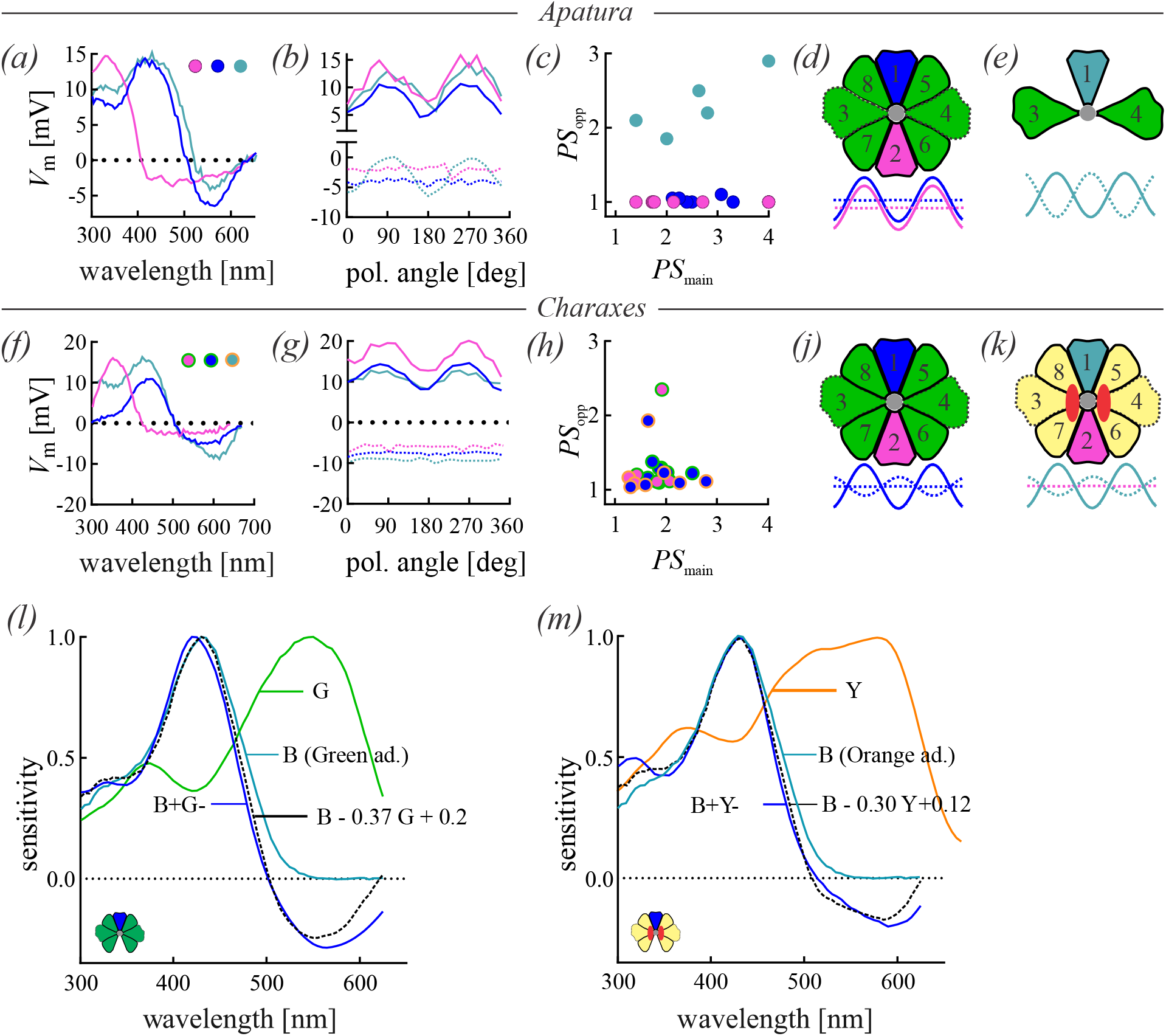
Opponent combinations U+G−, U+Y−, B+G− and B+Y− in *Apatura* and *Charaxes*. ***(a)*** Linear spectral sensitivity *S* of single U+G− and B+G− cells. (***b***) Responses of cells in (*a*) to flashes through a rotating polarizer; oscillation at positive and negative voltages indicates responses of the main unit (UV+, B+) and opponent (G−) cells. (***c***) PS of the opponent unit (G−) as a function of PS of the main unit (UV+, B+) in 13 UV+G− and 8 B+G− cells. ***(d)*** U+G− and B+G− with low PS of the opponent unit can be explained by the diagonal SVF R5-R8 or all six SVF R3-8 converging on a single postsynaptic R1,2 cell. The PS of the summed opponent inputs is cancelled out. ***(e)*** B+G− cells with high PS and orthogonal orientation of the opponent unit are explained by horizontal R3,4 converging on a single vertical R1,2 cell. In this case, R1,2 and R3,4 could form polarization-opponent pairs, suitable for the analysis of polarized light. (***f***) *S* of single U+G−, B+G− and B+Y− cells. (***g***) Responses of cells in ***(f)*** to flashes through a rotating polarizer; oscillation at positive and negative voltages indicates responses of the main (U+, B+) and opponent (G−, Y−) cells. (***h***) PS of opponent (G−, Y−) as a function of PS of main (U+, B+) cells. Scatterplot for 5 U+G−, 6 B+G− and 2 B+Y− cells. (***j, k***) U+G−, B+G− (*j*) and B+Y− (*k*) with low PS of the opponent unit are explained by presynaptic R3-8 converging on a single postsynaptic R1,2 cell. The PS of the summed input of the opponent cells is cancelled out. (***l, m***) Spectral sensitivities *S*_B+G−_ and *S*_B+Y−_ cells can be explained by linear subtraction of *S*_OPP_ of opponent cells from *S* of main cells.

#### Linear opponency model

Blue-sensitive cells in *Charaxes* have two different types of *S* curves in the opponent LW part. Maximal inhibition is either at ∼560 nm or at 590 nm in class B+G− and class B+Y− cells, respectively.

We tested whether their *S* curves could be explained by a linear subtraction of *S*_OPP_ from *S* of the cells with silenced opponent signals. We isolated the *S*_B_ of the blue units (teal line in (*l,m*), average of 7 cells) by selectively adapting their opponents with monochrome green or orange light, and subtracted linearly scaled S_G_ (*l*; average of 23 cells) and S_Y_ (*m*; average of 13 cells), respectively:

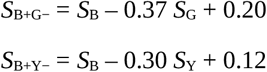

The numerical difference (dashed line in *m, l*) closely matched the experimentally measured *S*_B+G−_ and *S*_B+Y−_. The opponent signals in all but one B cells had PS∼1, meaning that in B+G− (*j*) and B+Y− (*k*), multiple G or Y cells converge on single B cells.

The opponency can be explained as a functionality of single ommatidia that contain only class G or only class Y R3-8 receptors. We extend by assuming that R3-8 cells from class G, and R3-8 cells from class Y R3-8 are present in ‘non-red’ and red-reflecting ommatidia, respectively.

### iv. Detailed analysis of stimulus response functions

**Figure S4.**
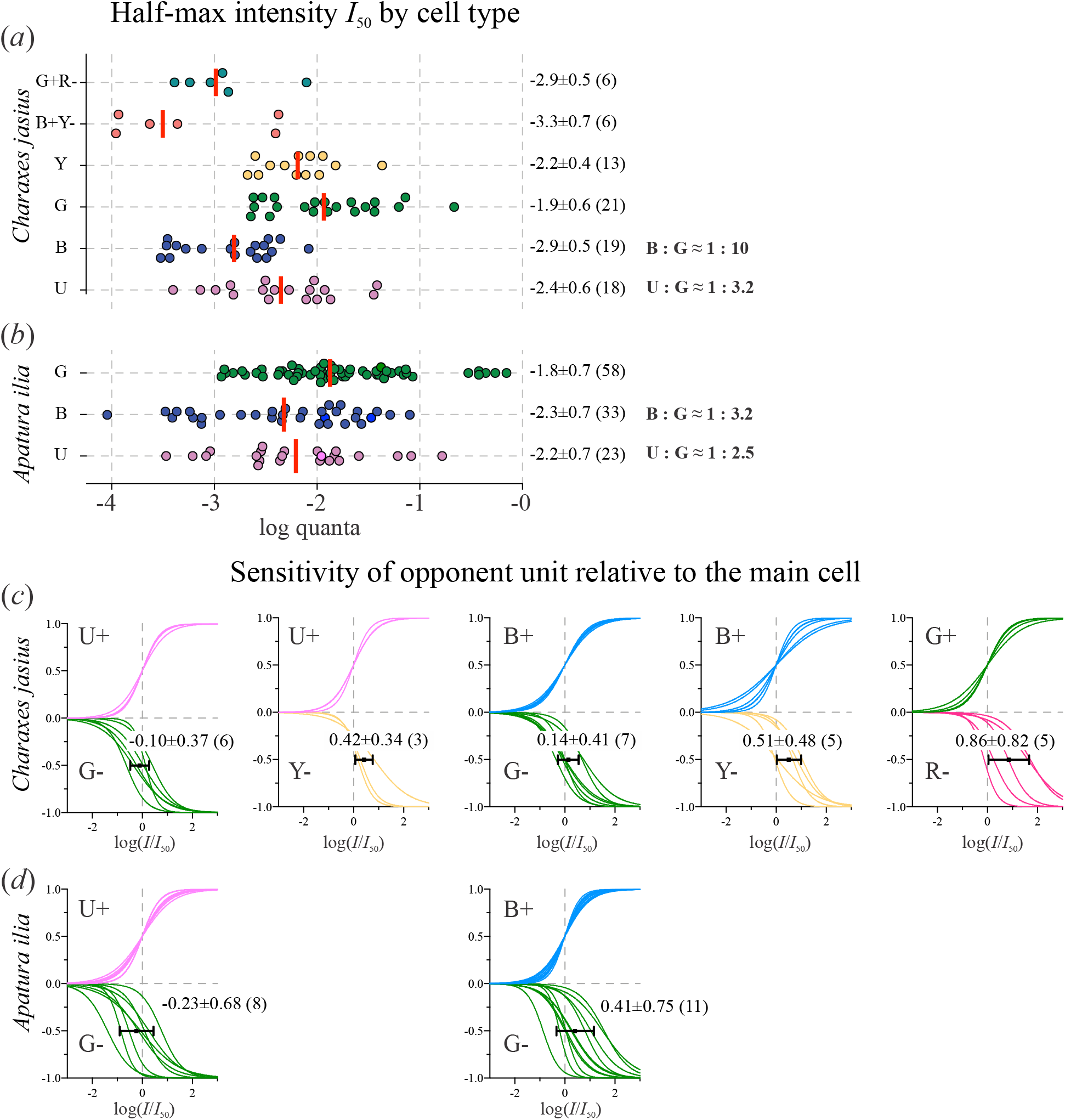
Sensitivity of the photoreceptors in *Charaxes* and *Apatura*. **(*a***,***b*)** Relative light intensity, evoking half-maximal response (*I*_50_) in the different spectral classes of photoreceptors in *Charaxes* (*a*) and *Apatura* (*b*). At log 0, flux is I ≈ 1.8×10^12^ photons cm^−2^ s^−1^. **(*c***,***d*)** Relative light intensity, evoking half-maximal (*I*_50_) opponent signals from the opponent units {G−, Y−, R−}, compared to the *I*_50_ of the main unit {U+, B+, G+} that were normalised to log 0. In both species, units G− in the class U+G− are slightly more sensitive (0.1∼0.2 log) than the main unit U+. In the other classes, the opponent units are less sensitive (∼0.1 log in B+G−, ∼0.9 log in G+R−).

### v. Anatomy of *Charaxes* compound eye

**Figure S5:**
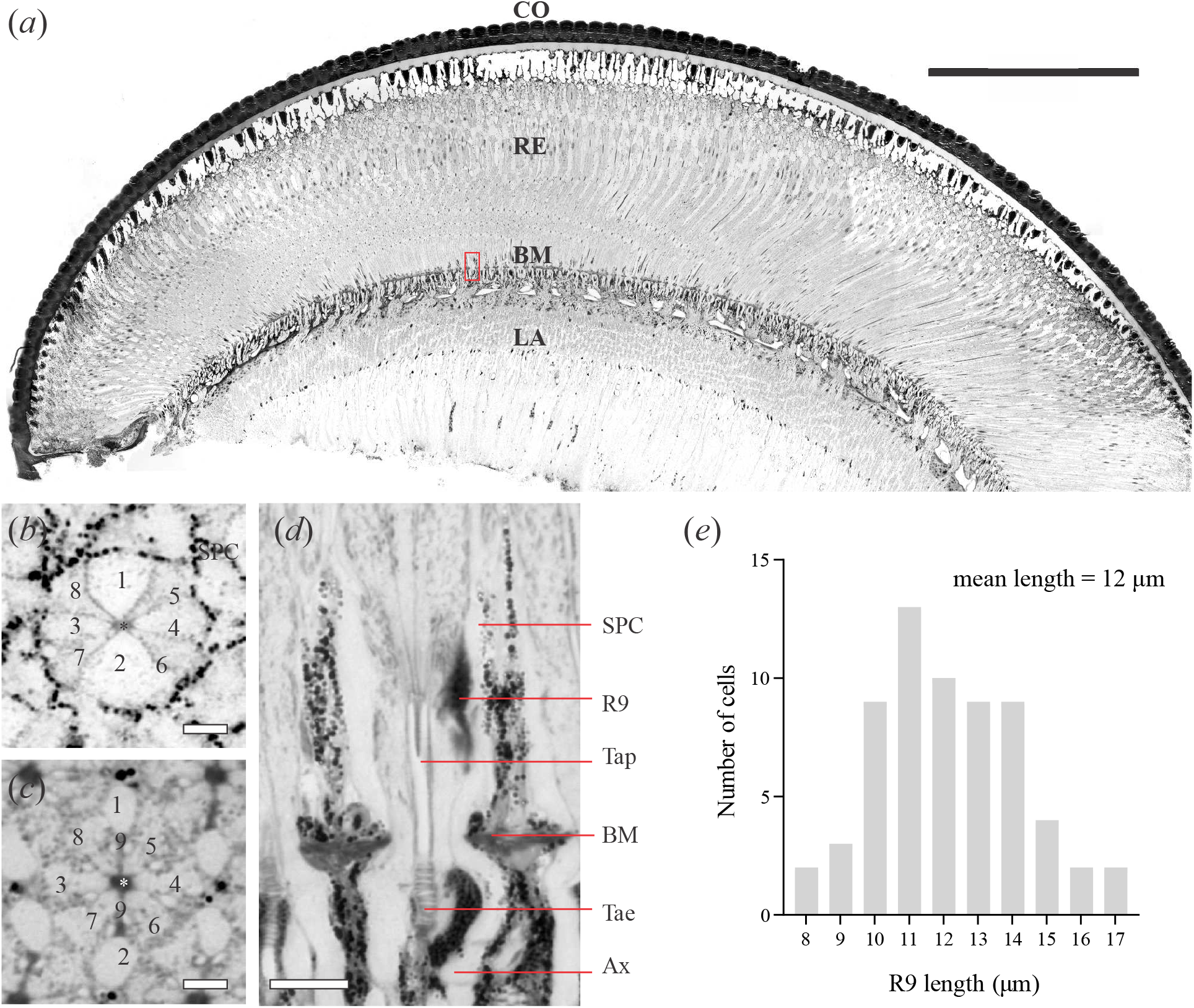
Light microscopical images of semithin sections of *Charaxes*. (***a***) Longitudinal section; **CO**, cornea; **RE**, retina; **BM**, basement membrane; **LA**, lamina; red rectangle magnified in (*d*). (***b***) Cross section at 214 μm. Receptor identity indicated by numbers; asterisk indicates rhabdom; **SPC**, secondary pigment cells. (***c***) Cross section at 478 μm. Receptor identity is indicated with *numbers*; *asterisk* **\*** indicates the rhabdom. At this level, R1&2 are axons and do not contribute to the rhabdom; R9 is dark and bilobed. (***d***) Longitudinal section, indicated with red rectangle in (*a*). **SPC**, secondary pigment cells; **R9**, basal receptor R9; **Tap**, tracheolar tapetum; **BM**, basement membrane; **Tae**, taenidia; **Ax**, receptor axon. (***e***) Length of the basal receptor R9 in 63 ommatidia. *Scale bar*, (*a*) 500 μm, (*b,c*) 5 μm, (*d*) 20 μm.

### vi. Spectral and polarisation properties of photoreceptors in *Danaus, Morpho, Archaeoprepona*

**Figure S6.**
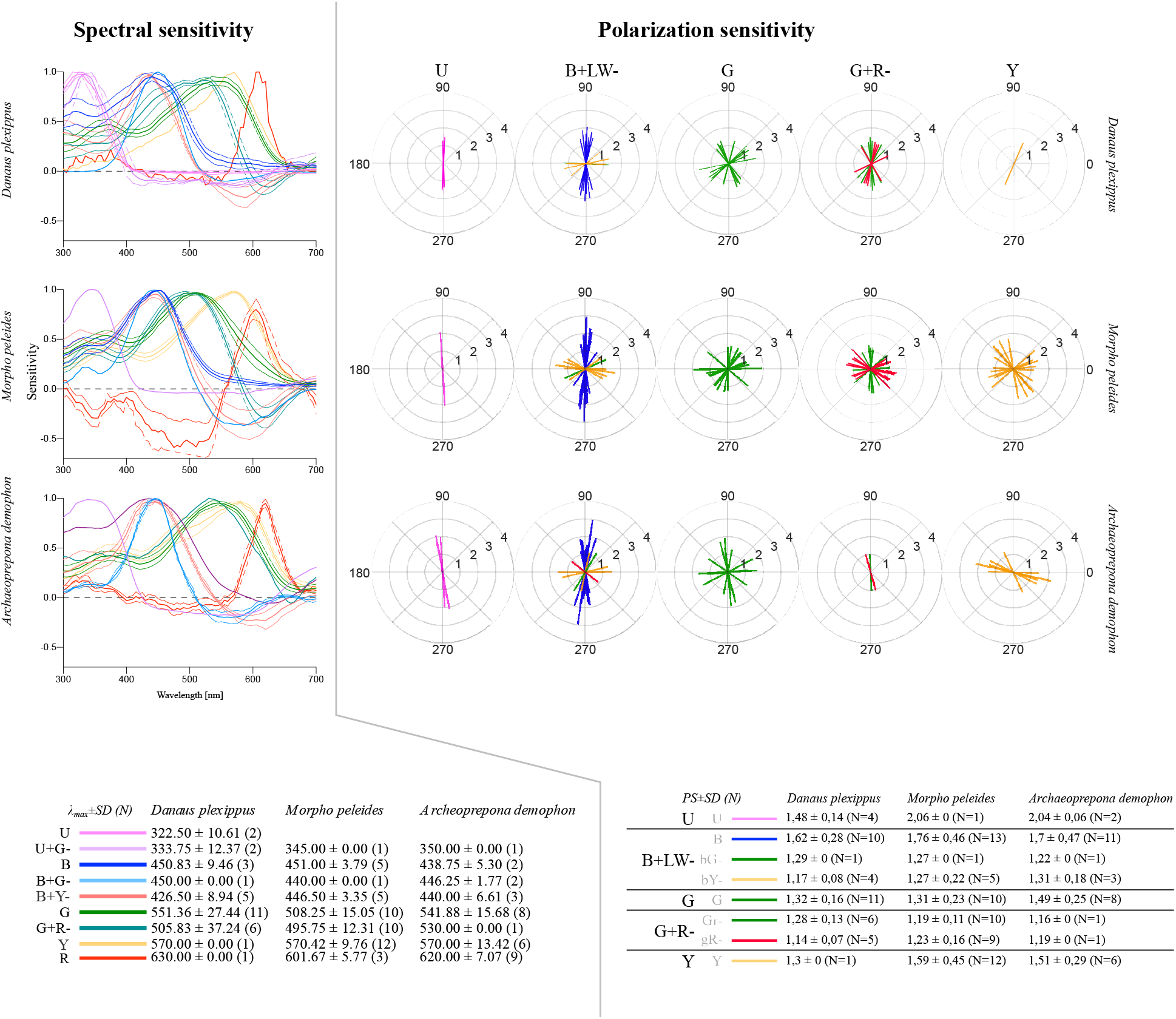
Spectral and polarization sensitivity in the monarch, blue morpho and prepona. These species all have an expanded lattice mosaic, the eyeshine exhibits red ommatidia. Similar as in *Charaxes*, their retina contains the basic set of photoreceptor classes (U+G−, B+G−, G_SVF_), and the expanded set with classes (B+Y−, Y, G+R−). The class U+Y− was not found, probably due to the small number of impaled class U photoreceptors. Receptor sensitivity maxima (λ_max_ in nm), polarization sensitivities are indicated in the tables at the bottom. Counts of analysed cells for each spectral class are in parenthesis. For eyeshine images, see our recent article (Belušič et al. 2021, reference [7]).

